# Exogenous DNA upregulates DUOX2 expression and function in human pancreatic cancer cells by activating the cGAS-STING signaling pathway

**DOI:** 10.1101/2022.05.14.491678

**Authors:** Stephen L. Wang, Yongzhong Wu, Mariam Konaté, Jiamo Lu, Smitha Antony, Jennifer L. Meitzler, Guojian Jiang, Iris Dahan, Agnes Juhasz, Becky Diebold, Krishnendu Roy, James H. Doroshow

## Abstract

Pro-inflammatory cytokines upregulate the expression of the H_2_O_2_-producing NADPH oxidase dual oxidase 2 (DUOX2) which, when elevated, adversely affects survival from pancreatic ductal adenocarcinoma (PDAC). Because the cGAS-STING pathway is known to initiate pro-inflammatory cytokine expression following uptake of exogenous DNA, we examined whether activation of cGAS-STING could play a role in the generation of reactive oxygen species by PDAC cells. Here, we found that a variety of exogenous DNA species markedly increased the production of cGAMP, the phosphorylation of TBK1 and IRF3, and the translocation of phosphorylated IRF3 into the nucleus, leading to a significant, IRF3-dependent enhancement of DUOX2 expression, and a significant flux of H_2_O_2_ in PDAC cells. However, unlike the canonical cGAS-STING pathway, DNA-related DUOX2 upregulation was not mediated by NF-κB; and although exogenous IFN-β significantly increased Stat1/2-associated DUOX2 expression, intracellular IFN-β signaling that followed cGAMP or DNA exposure did not itself increase DUOX2 levels. Finally, DUOX2 upregulation subsequent to cGAS-STING activation was accompanied by the enhanced, normoxic expression of HIF-1α as well as DNA double strand cleavage, suggesting that cGAS-STING signaling may support the development of an oxidative, pro-angiogenic microenvironment that could contribute to the inflammation-related genetic instability of pancreatic cancer.

## INTRODUCTION

Dual oxidase 2 (DUOX2) is an NADPH oxidase family member that plays an important role in mediating innate immunity at mucosal surfaces. Reactive oxygen species (ROS) produced by DUOX2 contribute to chronic inflammation-related tissue injury as well as angiogenesis, and can support the growth of epithelial malignancies [1-3]. DUOX2 expression is significantly increased in patients with chronic pancreatitis; furthermore, patients with repetitive bouts of pancreatic inflammation are predisposed to develop pancreatic ductal adenocarcinoma (PDAC), suggesting that DUOX2-mediated ROS could play a role in pancreatic carcinogenesis [4,5]. Our recent studies focusing on the control of DUOX2 expression have revealed that pro-inflammatory cytokines, including IFN-γ, IL-4, and IL-17A, upregulate DUOX2 expression in pancreatic cancer cells, producing oxidative DNA damage and DNA double strand breaks that could contribute to the pathogenesis of PDAC [4,6,7].

The cGAS-STING (cyclic GMP-AMP Synthase [cGAS]; Stimulator of Interferon Genes [STING]) signaling axis has been shown to play a vital role in innate immunity, protecting the host from viral infection. This signaling axis has also been demonstrated to both promote cancer progression and oncogenesis, as well as to enhance antitumor immunity [8-11]. Intratumoral injection of cyclic GMP-AMP (cGAMP) in murine cancer models produces an accumulation of macrophages in the microenvironment of various malignancies such as breast cancer, melanoma, and colon cancer which subsequently leads to the recruitment of CD8^+^ T cells that secrete a variety of pro-inflammatory cytokines (TNF-α, IFN-β) [11]. However, activation of the cGAS-STING pathway has also been demonstrated to stimulate carcinogenesis, in part by supporting an immunosuppressive and pro-metastatic microenvironment [12] as well as suppressing DNA repair [9].

The cytosolic DNA sensor cGAS detects and binds double-stranded DNA (dsDNA) that is ∼90 bp in length and longer [13] in a sequence-independent fashion. It then catalyzes the formation of cGAMP from GTP and ATP [14,15]. cGAMP in turn binds to the ER-bound protein STING which translocates to the Golgi and recruits Tank-binding kinase 1 (TBK1) and Interferon regulatory factor 3 (IRF3). Subsequently, TBK1 phosphorylates itself as well as IRF3 [16]. Phosphorylated IRF3 can then translocate into the nucleus along with NF-κB to promote the transcription of Type I Interferons (IFN) [14,17,18].

cGAS-STING appears to be involved in both the initiation and progression phases of PDAC [19,20]. Activation of STING signaling and enhancement of pancreatic inflammation was demonstrated in a murine model of pancreatitis. Zhao and colleagues found that DNA released by necrotic pancreatic acinar cells was taken up by phagocytes in the microenvironment and activated STING signaling and production of Type 1 IFN [21].

The importance of the pancreatic tumor microbiome for oncogenesis, cancer progression, and patient outcomes has recently been demonstrated [22,23]. Enrichment of certain microbes in the pancreatic tumor microbiome can contribute to pro-cancer phenotypes, depending on the type and genus of microbe involved. In murine models, gram-negative bacteria can traverse the intestine to reside in the normal pancreas; and in certain patients with PDAC, translocation of gram-negative *Proteobacteria* from the gut to the pancreas drives immune suppression and disease progression [22].

The mechanisms by which microbes or DNA released from necrotic cells into the microenvironment affect pancreatic cancer cells at the molecular level remain elusive despite increasing knowledge about the influential role that the pancreatic tumor microbiome plays in the severity and outcome of PDAC [24]. In this study, we report that uptake of exogenous DNA into the cytosol induced measurable levels of cGAMP synthesis and significantly enhanced DUOX2 expression at both the mRNA and protein levels in a panel of human PDAC cell lines following the activation of cGAS-STING signaling. Importantly, exposure to exogenous cGAMP as well as siRNA knockdown of cGAS confirmed the requirement for cGAS expression and enzymatic function in DNA-mediated enhancement of DUOX2 expression. Notably, cGAS-STING-mediated enhancement of DUOX2 expression was also associated with an increase in normoxic HIF-1α expression, H_2_O_2_ formation, and the production of DNA double strand breaks in PDAC cells. Consistent with the known deregulation of STING signaling in colon cancer [25], DUOX2 expression was not enhanced by exogenous DNA in human colon cancer cell lines, suggesting that the crosstalk between cGAS-STING signaling and DUOX2 is context dependent for tumors of the gastrointestinal tract.

In summary, these data suggest that extracellular DNA of mammalian or bacterial origin, by activating the cGAS-STING pathway, could support a DUOX2-induced, H_2_O_2_-mediated pro-inflammatory milieu that produces DNA double strand breaks which may contribute to the pathogenesis of PDAC.

## RESULTS

### Exogenous DNA activates cGAS-STING signaling and enhances DUOX2 expression in human pancreatic cancer cells

Uptake of DNA into cell cytosol from either necrotic cell debris, the formation of micronuclei due to DNA damage, or pathogens is common in areas of inflammatory tissue injury or abnormal tissue growth, such as in the tumor microenvironment [24]. Because recent studies from our laboratory have demonstrated the potential of pro-inflammatory cytokines to enhance oxidative stress in pancreatic ductal adenocarcinoma (PDAC) cells [7], we examined the effects of exogenous DNA on the NADPH oxidase family member that is prevalent in PDAC cells, DUOX2. We first evaluated the expression of STING and cGAS in PDAC cell lines that we have previously examined for their response to cytokines; we found that BxPC-3 and CFPAC-1 cells express both STING and cGAS protein in amounts that are easily demonstrable, whereas STING protein expression is limited in the AsPC-1 line (Fig. 1A). DUOX2 mRNA expression is significantly enhanced in BxPC-3, CFPAC-1, and HTB134 cells 48 h following transfection of exogenous DNA into cytosol (Fig. 1B, C, D). For the CFPAC-1 and HTB134 cell lines, the effect of transfected DNA plasmids is similar to the effects of exposure to the pro-inflammatory cytokines IL-17A or IL-4 for 24 h, respectively. As demonstrated in Fig. 1E, DNA plasmid increases the protein expression of DUOX 48 h after transfection; upregulation of DUOX occurs concomitant with the phosphorylation of TBK1 and IRF3. The enhanced DUOX level is also associated with increased expression of HIF-1α and the presence of DNA double strand scission in BxPC-3 cells as measured by the production of γH2AX. We found that total IRF3 and STING expression were diminished following plasmid transfection, an observation that is consistent observation that siRNA knockdown with previous studies describing the degradation of IRF3 after sustained activation of cGAS-STING signaling [21,26-28]. On the other hand, an increase in DUOX expression produced by exposure to IFN-γ for 24 h was, as expected, accompanied by a strong Stat1 phosphorylation signal without activation of TBK1 or IRF3.

**Fig. 1.**
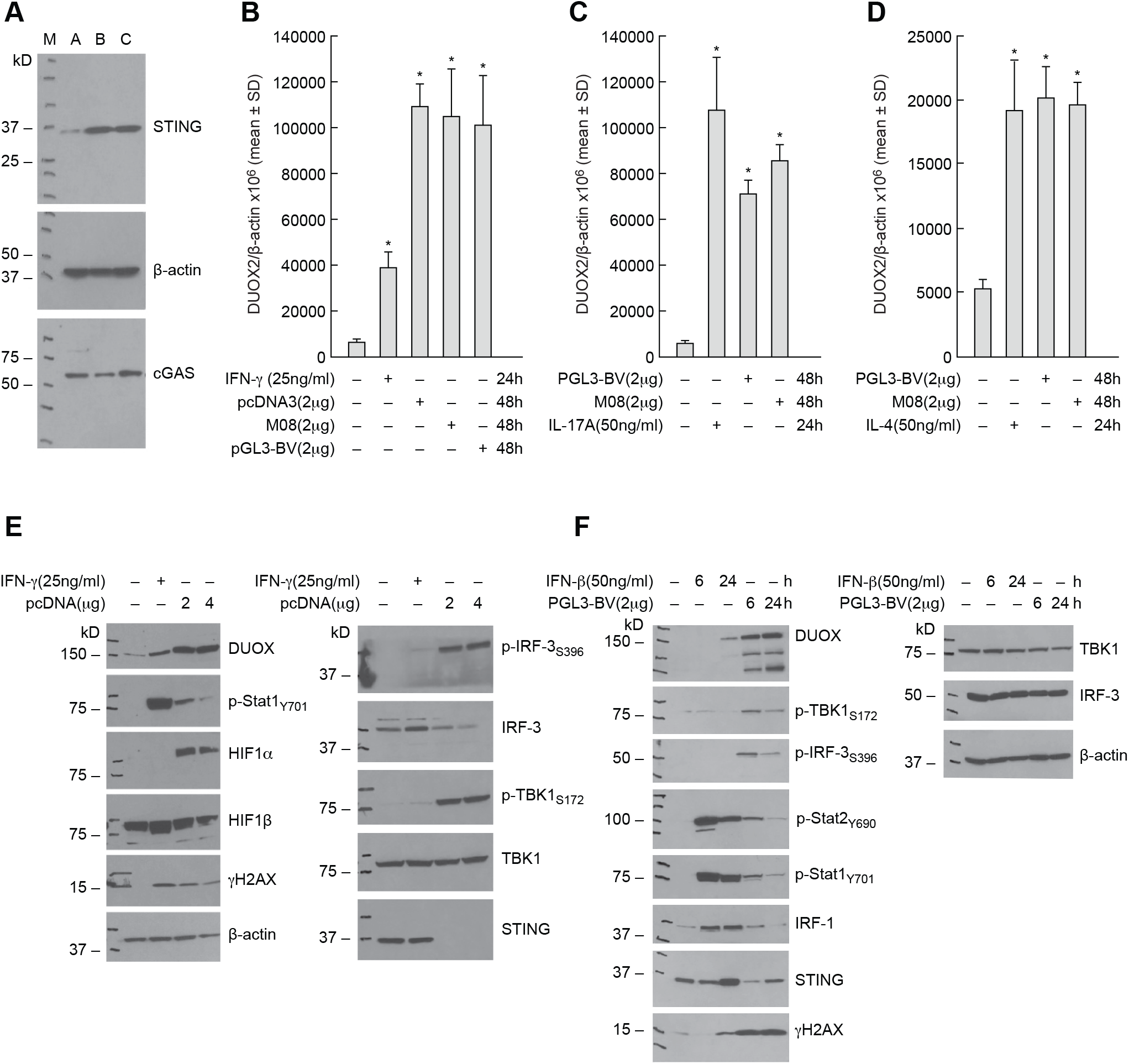
Introduction of exogenous DNA activates cGAS-STING signaling and enhances the expression of DUOX2 in human pancreatic cancer cells. **A** The protein expression levels of STING and cGAS were examined in three human pancreatic cancer cell lines: A: AsPC-1; B: BxPC-3; and C: CFPAC-1. **B** The effects of three different DNA plasmids on DUOX2 mRNA expression in BxPC-3 cells 48 h following transfection were examined by RT-PCR and compared to the effect of IFN-γ on DUOX2 expression used here as a positive control [4]. \**P* < 0.05. **C** DUOX2 expression in CFPAC-1 cells was determined 48 h after transfection of two DNA plasmids and compared to the effect of exposing these cells to IL-17A for 24 h [7]. \**P* < 0.05. **D** DUOX2 expression was measured by RT-PCR in HTB134 human pancreatic cancer cells 48 h following plasmid transfection or after exposure to 50 ng/ml IL-4 for 24 h [7]. \**P* < 0.05. **E** DUOX expression, cell signaling, DNA damage, and cGAS-STING activation were examined in BxPC-3 cells by Western blot following exposure to pcDNA plasmid (48 h following transfection) compared to the same cell line treated for 24 h with 25 ng/ml of IFN-γ. **F** CFPAC-1 cells were evaluated in these experiments to compare the time course of DUOX expression and activation of the cGAS-STING pathway following transfection of a DNA plasmid or exposure to IFN-β in complete media. All of the experiments shown here were repeated at least in triplicate.

To confirm these results, we evaluated the time-dependent activation of cGAS-STING signaling in a second PDAC cell line, CFPAC-1 (Fig. 1F). In these experiments, plasmid-related activation of the cGAS-STING pathway was demonstrable as early as 6 h following transfection, as shown by phosphorylation of TBK1 and IRF3; DUOX expression was also increased 6 h following plasmid exposure and was accompanied by evidence of enhanced DNA double strand breakage. We also found that treatment with IFN-β for 24 h upregulated DUOX expression as well as phosphorylation of Stat1 and Stat2 and the expression of IRF1 and γH2AX.

To evaluate the specificity of our results with human PDAC cells, we examined the effect of exogenous DNA on NADPH oxidase expression in human colon cancer cell lines (Supplementary Fig. S1). Transfection of plasmid DNA had no significant effect on either NOX1 or DUOX2 expression in the Ls513 line, nor on DUOX2 mRNA expression in either T84 or HT-29 colon cancer cells. However, in the same cell lines, proinflammatory cytokine exposure significantly increased DUOX2 mRNA levels (Supplementary Fig. S1A, B, C). In a previous study, double stranded DNA was found to produce a limited effect on type I IFN production by the HT-29 cell line [25]. Taken together, these data suggest that the presence of intracellular DNA activates cGAS-STING signaling and induces DUOX2 expression primarily in pancreatic rather than colon cancer cells.

### Concentration- and time-dependent enhancement of DUOX expression following DNA transfection is associated with increased H_2_O_2_ production by PDAC cells

Next, we examined the effect of DNA concentration and time following transfection on the expression of DUOX in PDAC cells. A plasmid level as low as 500 ng DNA significantly increased DUOX2 mRNA expression 48 h after transfection of BxPC-3 cells (*P* < 0.01, left panel, Fig. 2A). The same amount of DNA increased TBK1 phosphorylation and DUOX protein expression (middle panel, Fig. 2A). DNA-dependent upregulation of DUOX2 mRNA expression was significantly increased as early as 3 h following transfection in CFPAC-1 cells (*P* < 0.01, right panel, Fig. 2A). Transfection reagent alone in the absence of DNA produced no effect on the expression of DUOX2 (Fig. 2A, left and right panel). The upregulation of DUOX2 in BxPC-3 cells by DNA leads to the expression of a fully functional NADPH oxidase as shown in Fig. 2B. Forty-eight hours following plasmid transfection, BxPC-3 cells produce significantly higher levels of extracellular H_2_O_2_ (compared to solvent-treated control cells) as measured by the Amplex Red^®^ assay, *P* < 0.05. For comparative purposes, the rate of H_2_O_2_ production by BxPC-3 cells exposed for 24 h to IL-4 is also shown; results are consistent with the effect of IL-4 on DUOX2 expression and H_2_O_2_ production by BxPC-3 cells that we have demonstrated previously [7].

**Fig. 2.**
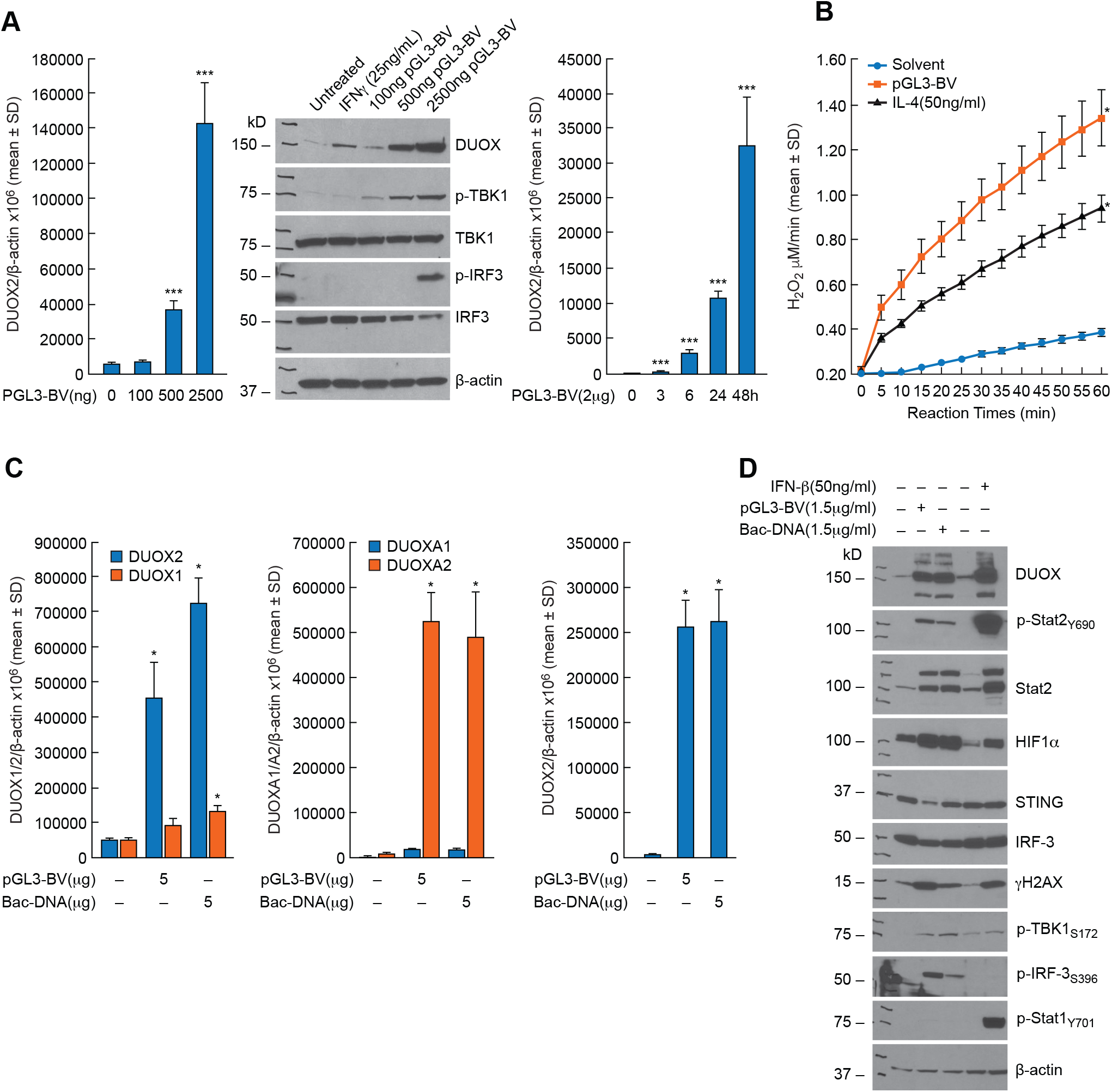
Concentration- and time-dependent enhancement of DUOX expression by plasmid DNA leads to significantly increased H_2_O_2_ production in PDAC cells while exogenous bacterial DNA is as effective as plasmid DNA in stimulating cGAS-STING signaling. **A** 48-h following transfection of PGL3-BV plasmid into BxPC-3 cells DUOX expression is significantly increased in a concentration-dependent fashion (left panel); enhanced DUOX expression 48 h following plasmid transfection is accompanied by phosphorylation of TBK1 and IRF3 (middle panel). Time course for plasmid-enhanced DUOX2 expression in CFPAC-1 cells propagated in complete media is shown in the right panel. \*\*\**P* < 0.01. **B** Time-dependent H_2_O_2_ production by BxPC-3 cells was measured using the Amplex Red^®^ assay 48 h following transfection of 2µg of pGL3-BV plasmid; the rate of H_2_O_2_ formation was compared to that of solvent treated cells and to cells exposed to IL-4 for 24 h, a treatment that has been shown previously to enhance the expression of functional DUOX2. H_2_O_2_ production was measured in the presence of ionomycin (1 µM). * *P* < 0.05. **C** In the left and center panels, the expression of DUOX1 and DUOX2 as well as DUOXA1 and DUOXA2 were determined 48 h following transfection of a DNA plasmid or E. coli DNA (Bac-DNA) into BxPC-3 cells. \**P* < 0.05. The right panel demonstrates DUOX2 mRNA expression 48 h following transfection of either plasmid or E. coli DNA into CFPAC-1 cells. \**P* < 0.05. **D** For BxPC-3 cells, the upregulation of DUOX and the downstream effects of increased DUOX expression, including activation of HIF-1α and DNA double strand scission measured by γH2AX, were similar 48 h following transfection with either plasmid or E. coli DNA. The effects of a 24 h IFN-β exposure on DUOX expression, DNA damage, and interferon-related signaling pathways are also shown. All experimental results shown are the result of at least three independent experiments.

Because of the recent demonstration that gram-negative bacteria are found in both human and murine pancreatic cancers [22], we compared the effect of plasmid DNA to that of E. coli for the ability to induce DUOX2 expression in PDAC cell lines (Fig. 2C). E. coli DNA significantly increased DUOX2 and DUOXA2 levels in BxPC-3 cells to the same degree as plasmid DNA, *P* < 0.05; DUOX1 mRNA expression was also significantly increased but to a much smaller degree. In CFPAC-1 cells, DUOX2 expression was enhanced similarly by both plasmid and E. coli DNA. The western blot shown in Fig. 2D confirms that transfection of bacterial DNA into BxPC-3 cells activates the cGAS-STING pathway and enhances the expression of DUOX, HIF-1α, and γH2AX. Finally, in contrast to our findings for DUOX1 and DUOX2, we observed that transfection of either pGL3-BV plasmid or E. coli DNA into BxPC-3 cells did not increase the mRNA expression of either NOX1 or NOX4 (data not shown).

### Role of specific components of the cGAS-STING pathway in the upregulation of DUOX2 expression by exogenous DNA

Using siRNAs against cGAS, we found that the enhanced expression of DUOX2 following plasmid transfection could be significantly decreased in BxPC-3 cells when cGAS expression was diminished (Fig. 3A). cGAS siRNA blocked plasmid-stimulated DUOX protein expression, phosphorylation of TBK1 and IRF3, as well as expression of cGAS itself in these cells (Fig. 3B). Since cGAS binds to and is activated by double stranded DNA to produce cGAMP, we examined by ELISA whether pGL3-BV plasmid altered cGAMP levels in a concentration-dependent fashion 48 h following transfection. A positive linear relationship could be demonstrated between intracellular cGAMP levels and plasmid DNA, with an R^2^ of 0.94 for the BxPC-3 line (Fig. 3C). We also found using 2 µg of pGL3-BV DNA that the time course of cGAMP production following plasmid transfection rose linearly to ≈ 125 pg/ml for the first 8 h and then began to plateau for the subsequent 40 h of observation (data not shown). Because of the effect of transfected DNA on intracellular cGAMP levels in BxPC-3 cells, we examined whether exposure to extracellular cGAMP, which can be imported by tumor cells [29,30], altered DUOX2 mRNA expression. As shown in Fig. 3D, exposure of BxPC-3 cells for 24 h to extracellular cGAMP at a concentration of 25 µg/ml significantly increased DUOX2 mRNA expression, *P* < 0.05. This effect was time dependent, reaching significance following a 6 h exposure to 25 µg/ml cGAMP, left panel of Fig. 3E; cGAMP exposure also significantly increased IFN-β expression in BxPC-3 cells within 3 h, right panel of Fig. 3E. The time course for cGAMP-enhanced cGAS-STING signaling is shown in Fig. 3F; while increased DUOX protein expression is observed 24 h following cGAMP exposure, activation of TBK1 and IRF3 occur as early as 1 h following the addition of cGAMP, and evidence of IFN-β-related signal transduction (phosphorylation of Stat1 and Stat2 and increased expression of IRF9) can be demonstrated within 3 h. These experiments suggest that the cGAS-STING pathway, including upregulation of IFN-β-related signaling, is activated following the engagement/activation of cGAS by double stranded DNA in BxPC-3 cells. However, as shown in Fig. 3G, when examined concurrently, both the time course and the degree of IFN-β-related Stat phosphorylation differ when the effect of the exogenous type I interferon is compared to cGAMP treatment; IFN-β activates Stat1/2 within 1 h of cytokine exposure, and activation lasts for at least 24 h; whereas, treatment with cGAMP appears to activate Stat signaling to a lesser degree and for a shorter duration.

**Fig. 3.**
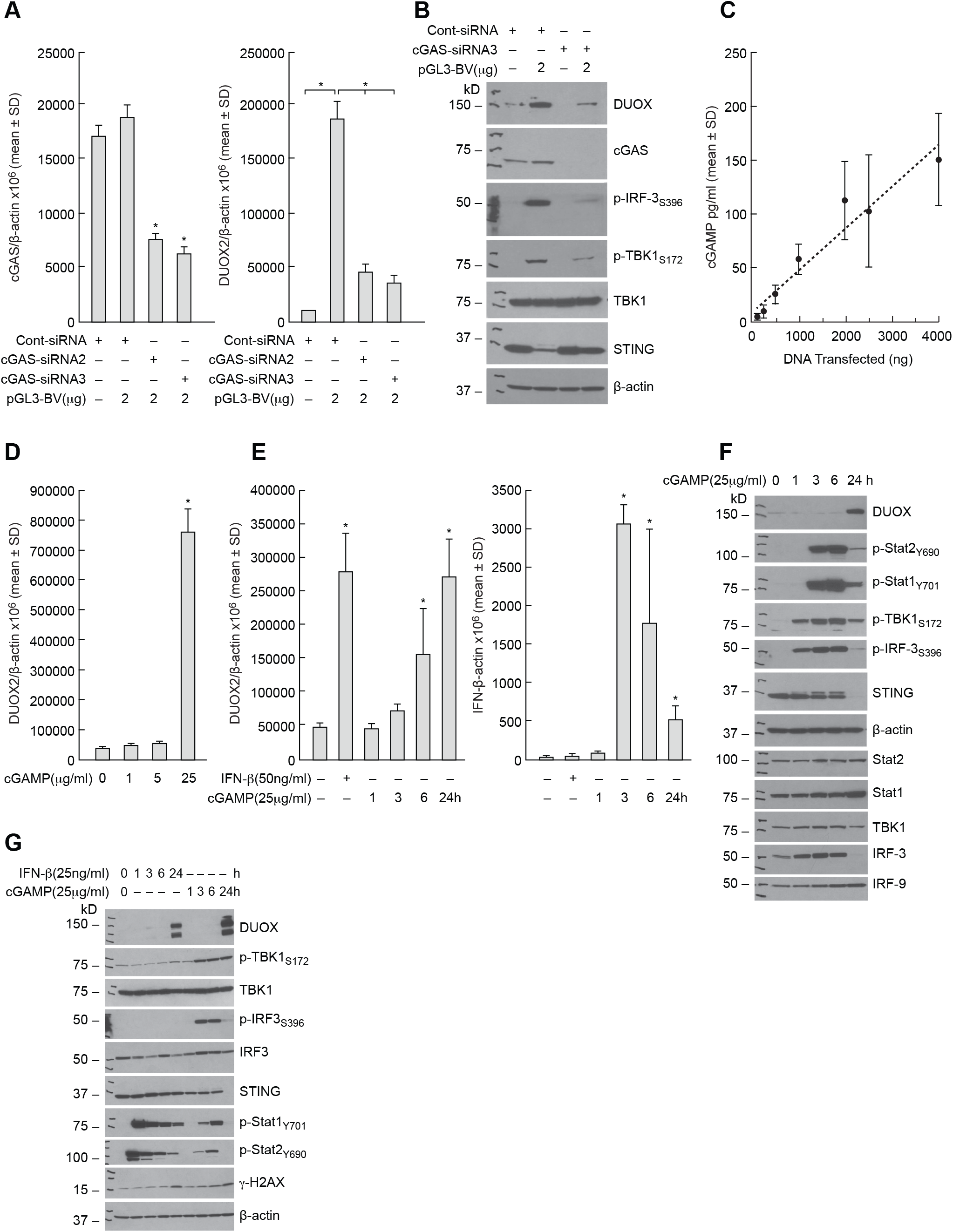
Role of the cGAS-STING pathway in the enhancement of DUOX2 expression by extracellular DNA in BxPC-3 pancreatic cancer cells. **A** Effect of intracellular cGAS levels (left panel) on expression of DUOX2 (right panel) 48 h following transfection of pGL3-BV plasmid examined using cGAS siRNA in BxPC-3 cells. **B** Evaluation of the role of cGAS in plasmid-enhanced DUOX protein expression and cell signaling in the BxPC-3 cell line. Western analysis was performed 48 h following plasmid and siRNA transfection. **C** DNA plasmid concentration-dependent increase in cellular cGAMP production by BxPC-3 cells. Tumor cells were transfected with increasing amounts of pGL3-BV DNA; 48 h following transfection, intracellular cGAMP was measured by ELISA. *P* < 0.05 for all DNA levels ≥ 500 ng. **D** Concentration-dependent increase in DUOX2 expression following 24 h exposure to extracellular cGAMP in BxPC-3 cells. \**P* < 0.05. **E** Left panel; time-dependent increase in cGAMP-related DUOX2 expression in BxPC-3 cells compared to the effect of 24 h IFN-β treatment on DUOX2 mRNA levels. Right panel; effect of cGAMP exposure time on expression of IFN-β mRNA. \**P* < 0.05. **F** Time course for cGAMP-related DUOX protein expression and cGAS-STING signaling. **G** Comparison of the time course of the effects of IFN-β and cGAMP on Stat and cGAS-STING signaling, DUOX expression, and DNA damage in BxPC-3 cells. All experiments shown in this figure were repeated a minimum of three times.

### Signal transduction downstream of activated cGAS-STING in PDAC cell lines

To examine cGAS-STING-dependent signaling in PDAC cells without transfecting DNA, we evaluated the effects of the STING agonist MSA-2 [31] on the events downstream of cGAS that may contribute to enhanced DUOX expression. MSA-2 treatment, similar to plasmid DNA, increases the expression of DUOX in a time-dependent fashion in BxPC-3 cells; increased DUOX expression is preceded by phosphorylation of TBK1 and IRF3 beginning 1 h following drug exposure (Fig. 4A). DNA double strand breakage occurs in concert with increased DUOX expression, while phosphorylation of Stat1 and Stat2, suggestive of IFN-β signaling, are demonstrable 6 h following initiation of MSA-2 exposure. On the other hand, for AsPC-1 cells, which demonstrate more modest baseline expression of STING compared to BxPC-3 cells (Fig. 1A and Fig. 4A), MSA-2 failed to activate IRF3 or Stat transcription factors and did not increase DUOX expression. Significant enhancement of DUOX2 expression by MSA-2 is concentration dependent in both BxPC-3 (Supplementary Fig. S2A) and CFPAC-1 (Supplementary Fig. S2B) cells; furthermore, the time course of MSA-2 activation of cGAS-STING signaling resembles that produced by exposure to cGAMP, differing only in the modest activation of Stat2 by the STING agonist (Supplementary Fig. S2C). Finally, because we previously demonstrated that dexamethasone co-treatment blunts pro-inflammatory cytokine-related upregulation of DUOX [32], we evaluated the effect of the glucocorticoid on signal transduction following cGAMP and MSA-2 exposure. Dexamethasone treatment partially blocks DUOX upregulation by either agent, as well as DNA double strand breakage, and cGAS-STING-mediated phosphorylation of IRF3 and TBK1 (Supplementary Fig. S2D).

**Fig. 4.**
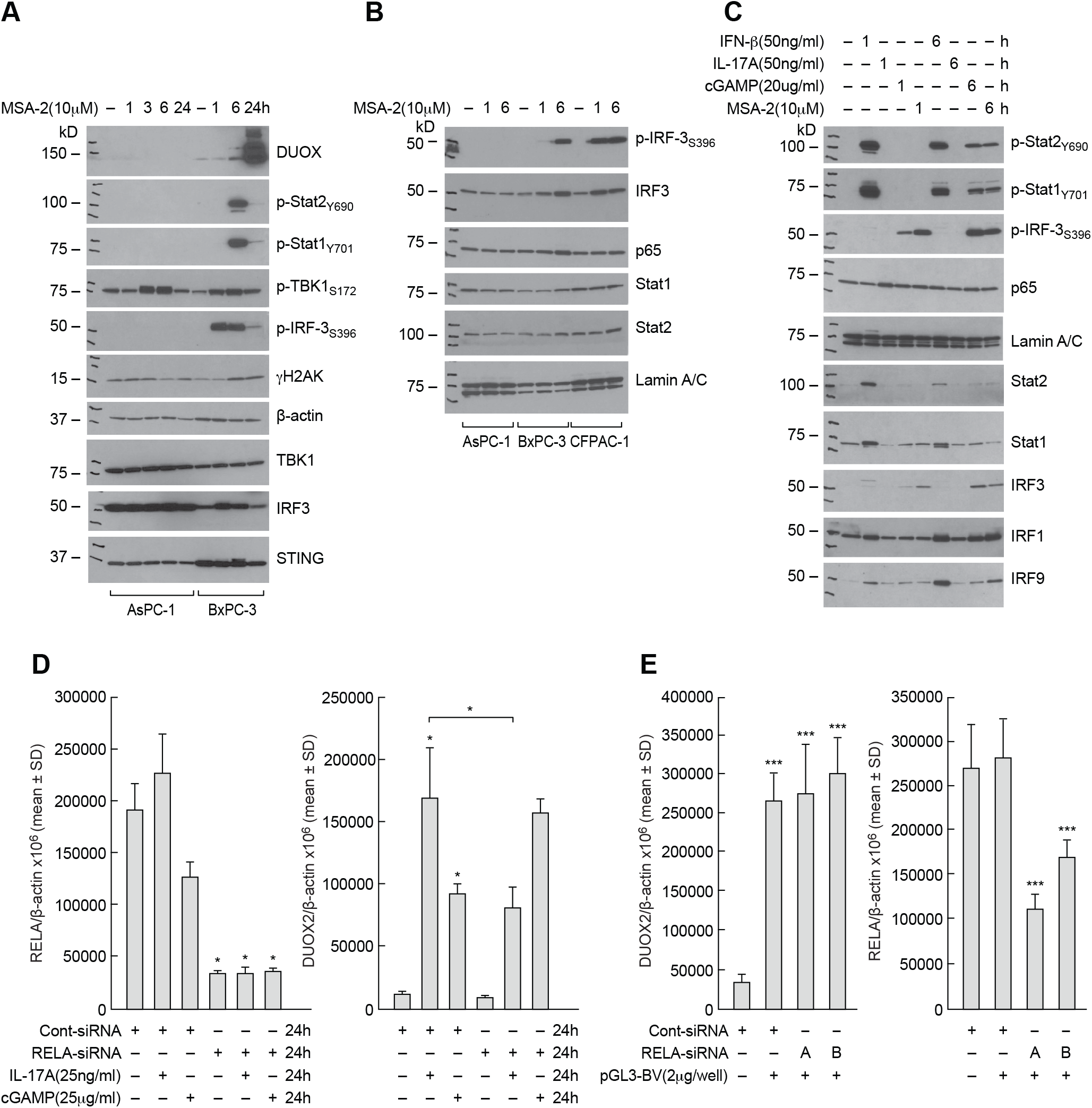
Activation of signaling pathways downstream of cGAS-STING in PDAC cell lines. **A** Comparison of cell signaling and DNA damage time course following exposure to the STING agonist MSA-2 (10 µM) in AsPC-1 and BxPC-3 cells. **B** Time course for cGAS-STING and cytokine nuclear signaling following exposure to 10 µM MSA-2 in AsPC-1, BxP-3, and CFPAC-1 tumor cells. **C** Comparison of cGAMP/MSA-2 induced cell signaling to that produced by IFN-β and IL-17A in BxPC-3 cells. Cells were untreated or exposed for 1 or 6 h to each of the compounds studied. **D** Effect of NF-κB signaling on cGAMP-related DUOX2 expression. The role of RELA expression in cGAMP- and IL-17A-related DUOX2 (right panel) and RELA (left panel) expression was examined in BxPC-3 cells using siRNA. In these experiments, control siRNA and RELA siRNA, where indicated, were transfected into BxPC-3 cells; 24 h following transfection, cells were propagated in serum free medium alone or with the addition of either IL-17A or cGAMP for another 24 h. \**P* < 0.05. **E** Role of NF-κB signaling in plasmid-related upregulation of DUOX2 expression in BxPC-3 cells evaluated using two different RELA siRNAs; left panel demonstrates effects of RELA siRNAs on DUOX2 expression 48 h after plasmid transfection, and the right panel shows the effect of the siRNAs on RELA expression itself. \*\*\**P* < 0.01. The results presented represent at least three independent experiments.

We next examined the nuclear translocation of IRF3 and phospho-IRF3, p65, Stat1 and Stat2 following MSA-2 treatment in PDAC cell lines (Fig. 4B). Nuclear translocation of phosphorylated IRF3 by 6 h was clear for both BxPC-3 and CFPAC-1 but not AsPC-1 cells. However, translocation of the NF-κB component p65 (RELA) following MSA-2 exposure was not prominent in any PDAC cell line. To broaden our evaluation of nuclear signaling, we compared the effects of IFN-β, IL-17A, cGAMP, and MSA-2 on pathways downstream of cGAS-STING in BxPC-3 cells (Fig. 4C). As expected, IFN-β activates Stat1 and Stat2 and increases the expression of IRF9. Furthermore, exposure to both cGAMP and MSA-2 leads to the nuclear translocation of phosphorylated IRF3. However, only IL-17A treatment modestly increases the expression of p65 in the nucleus.

To explore the role of NF-κB signaling further, the effect of RELA siRNA on DUOX expression was studied in BxPC-3 cells. Despite > 75% knockdown of RELA mRNA expression (Fig. 4D, left panel), siRNA treatment did not decrease cGAMP-mediated upregulation of DUOX2 (Fig. 4D, right panel). On the other hand, the increase in DUOX2 levels produced by IL-17A, that we have previously shown to be regulated, in part, by NF-κB [7], was significantly decreased by RELA siRNA, *P* < 0.05. RELA siRNAs also did not diminish the significantly enhanced DUOX2 expression that occurred 48 h following plasmid transfection (Fig. 4E).

### IRF3, but not Stat1 or Stat2, plays an important role in cGAS-STING-mediated enhancement of DUOX2 expression

To further elucidate the mechanism by which exogenous DNA mediates increased DUOX2 expression following activation of cGAS-STING signaling, we employed an siRNA knockdown strategy. Knockdown of IRF3 expression in BxPC-3 by ∼50% (Fig. 5A, right panel) significantly decreased DNA-mediated DUOX2 mRNA expression relative to a scrambled siRNA control (Fig. 5A, left panel). On the other hand, siRNA knockdown of IRF1 did not alter DNA-mediated DUOX2 expression (Fig. 5A, left and middle panels; Supplementary Fig. S3G). These results were confirmed for MSA-2-mediated, as well as plasmid-mediated, DUOX2 expression using three different IRF3 siRNAs (Fig. 5B and 5C). Experiments evaluating CFPAC-1 cells exposed to cGAMP demonstrated similar results, confirming the role of IRF3 in cGAS-STING-mediated DUOX2 expression (data not shown). At the protein level, knockdown of IRF-3 with two different siRNAs markedly diminished plasmid-enhanced DUOX and HIF-1α expression in the BxPC-3 cell line (Fig. 5D).

**Fig. 5.**
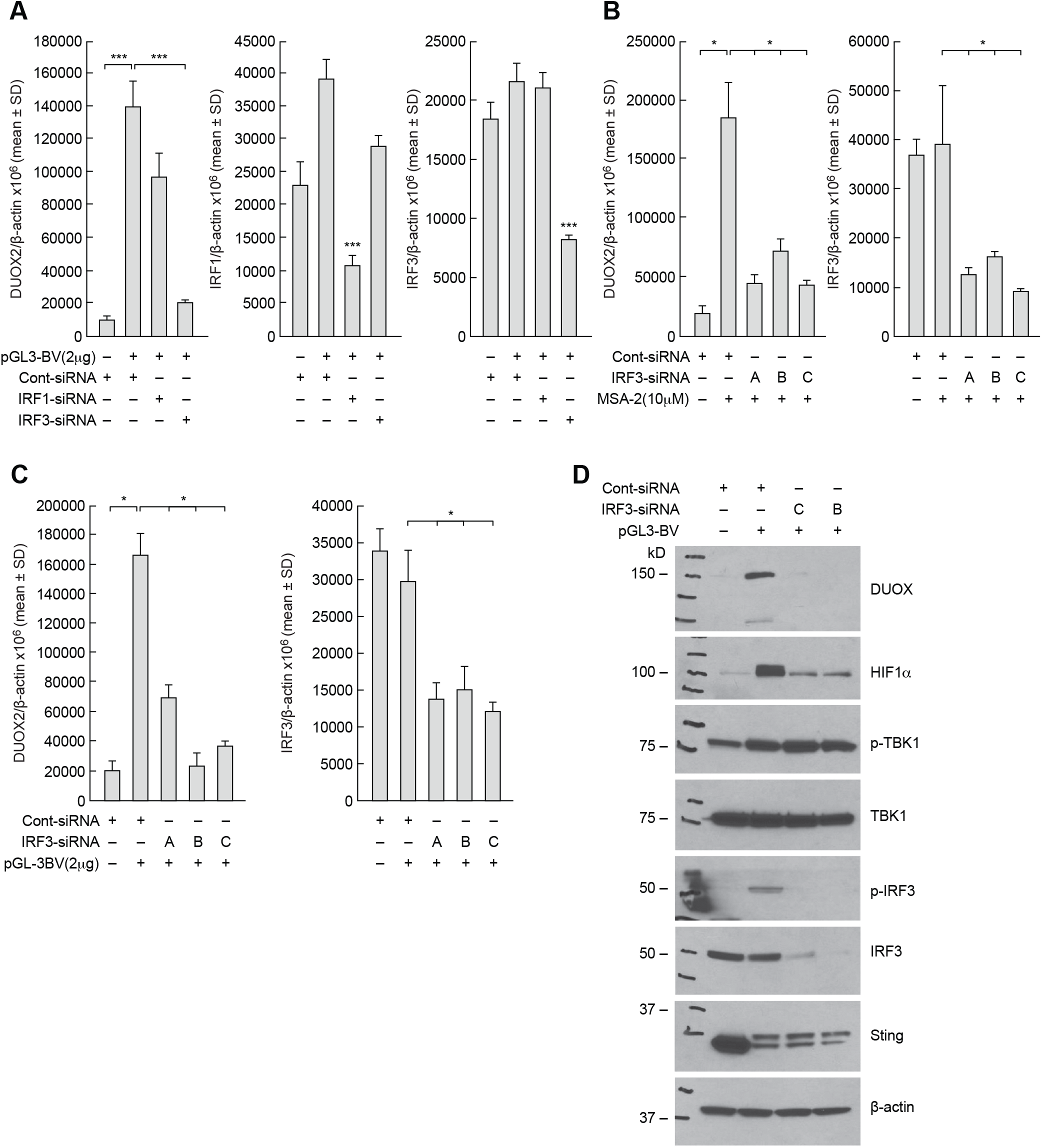
Role of IRF3 in the control of cGAS-STING-mediated enhancement of DUOX2 expression. **A** The contributions of IRF1 and IRF3 to plasmid-enhanced expression of DUOX2 measured using RT-PCR were examined in the BxPC-3 cell line using IRF1 or IRF3 siRNAs in the left panel, and on IRF1 or IRF3 expression levels in the middle and right panels, respectively. \*\*\**P* < 0.01. **B** In the left panel, the effect of IRF3 siRNAs on MSA-2-enhanced DUOX2 expression was determined for BxPC-3 cells; downregulation of IRF3 by siRNAs was examined in the right panel. \**P* < 0.05. **C** IRF-3 siRNAs block the upregulation of DUOX2 mRNA expression following plasmid transfection (left panel) and baseline IRF3 mRNA after pGL3-BV transfection (right panel) in BxPC-3 cells. \**P* < 0.05. **D** At the protein level, IRF3 siRNA diminishes the enhanced expression of DUOX by the pGL3-BV plasmid in BxPC-3 pancreatic cancer cells. These results are representative of three independent experiments.

Because we had demonstrated in these studies that DUOX2 expression is significantly enhanced when PDAC cells are treated with IFN-β (Fig. 3E), and that Stat signaling pathways that are downstream of IFN-β are activated by cGAMP and MSA-2 (Fig.4C), we examined the role of Stat signaling in DNA-mediated upregulation of DUOX2 (Supplementary Fig. S3). As shown in Supplementary Fig. S3A and Supplementary Fig. S3B, transfection of plasmid DNA into BxPC-3 cells significantly increases DUOX2 expression; however, knockdown of either Stat1 or Stat2 with siRNA does not alter DNA-enhanced upregulation of DUOX2 mRNA expression. Furthermore, while Stat2 knockdown blocks IFN-β-stimulated DUOX2 expression, at least in part, it does not inhibit MSA-2-related upregulation of DUOX2 (Supplementary Fig. S3C, left panel). These results were confirmed for IFN-β, using multiple Stat2 siRNAs (Supplementary Fig. S3D). Finally, the ineffectiveness of Stat2 knockdown on MSA-2-related upregulation of DUOX2 expression was confirmed for plasmid DNA- and cGAMP-enhancement of DUOX2 expression using multiple siRNAs (Supplementary Fig. S3E and Supplementary Fig. S3F).

## DISCUSSION

In a recent study, our laboratory demonstrated that high level DUOX2 expression is adversely correlated with survival in patients with PDAC and that T_H_2 and T_H_17 cytokines synergistically induce expression of DUOX2 in PDAC cell lines [7]. These experiments broadened the known range of pro-inflammatory cytokines, beyond IFN-γ and LPS, capable of enhancing DUOX2 expression and DUOX2-mediated H_2_O_2_ production in pancreatic cancer cells [33]. Because the cGAS-STING pathway plays an important role in the regulation of pro-inflammatory cytokine expression [34] and is known to be highly expressed in both human PDACs and in murine pancreatic cancer models [35], we examined whether a relationship might exist between cGAS-STING signaling and DUOX2 expression.

The demonstration that exogenous DNA from plasmids or bacterial sources significantly enhanced DUOX2 expression and function in a panel of PDAC cell lines in a concentration- and time-dependent fashion initially suggested that the canonical cGAS-STING pathway was operating in our studies (Fig. 6). Uptake of exogenous DNA led to the formation of cGAMP, the phosphorylation of TBK1 and IRF3, and the translocation of phosphorylated IRF3 to the nucleus of PDAC cells. These results were confirmed by exposure to exogenous cGAMP as well as by treatment with the STING agonist MSA-2 (Fig. 3D and 3E; Suppl. Fig. S2). Furthermore, cGAS siRNAs significantly diminished both cGAS expression and upregulation of DUOX2 by a DNA plasmid. We also found using multiple siRNAs that IRF3, and not IRF1, is necessary for DNA-mediated induction of DUOX2 expression (Fig. 5).

**Fig. 6.**
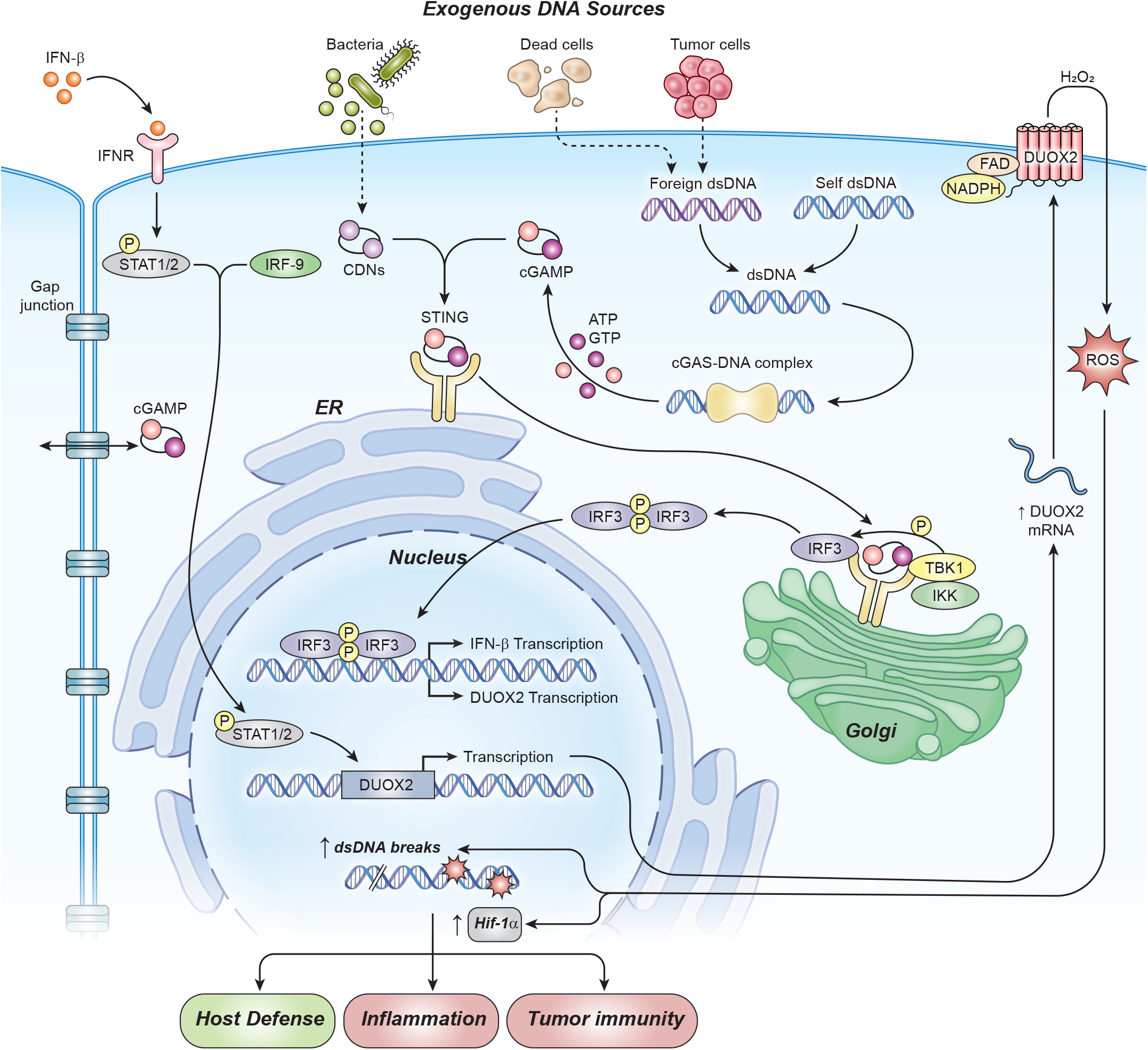
cGAS-STING-mediated enhancement of DUOX2 expression in PDAC cells. In this model, foreign DNA, from exogenous plasmids or from bacterial sources, when transferred into human PDAC cells activates cGAS-STING signaling. After binding DNA in the cytosol, cGAS catalyzes the formation of cGAMP from GTP and ATP. cGAMP in turn binds to the ER-bound protein STING which translocates to the Golgi and recruits Tank-binding kinase 1 (TBK1) and Interferon regulatory factor 3 (IRF3). The formation of this signaling complex allows TBK1 to phosphorylate IRF3 and auto-phosphorylate itself. For PDAC cells, in a non-canonical fashion, phosphorylated IRF3, but not NF-κB, appears to be responsible for enhancing the transcription of DUOX2 mRNA and the subsequent production of an enzymatically active oxidase that produces a flux of H_2_O_2_ capable of crossing cell membranes. Our experiments have also shown that extracellular cGAMP can be imported into PDAC cells, enhancing DUOX2 expression directly in the absence of extracellular DNA. Exogenous IFN-β signals downstream through Stat1/2 to increase DUOX2 protein expression. However, although exogenous IFN-β capably upregulates DUOX2, when the cytokine is generated intracellularly as a consequence of cGAS-STING signaling in PDAC cells, IFN-β signaling is limited and does not appear to contribute substantively to the expression of DUOX2. Reactive oxygen species generated by DUOX2 facilitate increased, normoxic expression of HIF-1α and DNA double strand cleavage that could sustain an oxidative, pro-inflammatory environment. Acutely, this might foster tumor immunity; however, chronic cGAS-STING-induced DUOX2 expression could promote DNA double strand breakage enhancing genetic instability. (Abbreviations used in the figure: DUOX2, dual oxidase 2; CDNs, cyclic dinucleotides; cGAS, cyclic GMP-AMP Synthase; STING, Stimulator of Interferon Genes; cGAMP, cyclic GMP-AMP; IFN-β, interferon beta; dsDNA, double stranded DNA; TBK1, Tank-binding kinase 1; IRF3, interferon regulatory factor 3; IRF9, Interferon regulatory factor 9; STAT1/2, Signal transducer and activator of transcription 1 or 2; ROS, reactive oxygen species; HIF-1α, Hypoxia-Inducible Factor 1; figure adapted from [37])

However, while exogenous DNA activated cGAS-STING signaling and generated cGAMP, activation of NF-κB with translocation to the nucleus was not demonstrated following cGAMP exposure; furthermore, although RELA siRNA partially inhibited the upregulation of DUOX2 by IL-17A (consistent with our prior experiments demonstrating the NF-κB-dependence of this effect), multiple RELA siRNAs did not alter either plasmid-or cGAMP-mediated enhancement of DUOX2 expression (Figs. 4D and 4E). The observation that siRNA knockdown of NF-κB does not appear to have a significant effect on DNA-mediated DUOX2 expression suggests that the mechanism that underlays the upregulation of DUOX2 by cGAS-STING diverges, in part, from the canonical pathway.

Moreover, we demonstrated for the first time that exposure to the pro-inflammatory cytokine IFN-β strongly upregulates DUOX2 expression in PDAC cells. DUOX2 upregulation by IFN-β was accompanied by increased expression of IRF-9, as well as phosphorylation of Stat1 and Stat2. This effect of IFN-β on DUOX2 expression might have been expected because of our previous demonstration that IFN-γ-mediated upregulation of DUOX2 is produced by Stat1 binding to the DUOX2 promoter [33]. However, despite the cGAMP-mediated increase in IFN-β expression that we observed (Fig. 3E), siRNAs against Stat1 and Stat2, although capable of blocking enhanced DUOX2 expression produced by IFN-β, did not inhibit MSA-2-or plasmid-mediated increases in DUOX2 levels (Suppl. Fig. S3). Thus, although the cGAS-STING pathway appears capable of regulating IFN-β transcription in PDAC cells, signaling by intrinsically-produced IFN-β does not appear to explain enhanced DUOX2 expression following the uptake of exogenous DNA.

In conclusion, our findings identify a novel crosstalk between the cGAS-STING immune sensing pathway and NADPH oxidase expression, both of which are implicated in mediating innate immunity at mucosal surfaces and in cancer progression. Since cGAS-STING-related enhancement of DUOX2 levels (and subsequent H_2_O_2_ formation) increases the normoxic expression of HIF-1α and promotes DNA double strand scission (as measured by γH2AX), our experiments suggest that peroxide-mediated DNA oxidation and double strand breaks occurring downstream of DUOX2 [7] might provide a feedback mechanism that could sustain cGAS-STING activation [36]. Such a process may contribute to oxidant-related pancreatic carcinogenesis stimulated by extracellular DNA present in a pro-inflammatory pancreatic microenvironment.

## MATERIALS AND METHODS

### Cell culture, antibodies, and reagents

All cell lines, culture conditions, antibodies, plasmids, primers, and reagents are described in Supplementary Methods.

### Western analysis

Tumor cells (4 × 10^6^) were plated in 100 mm dishes (Corning) and harvested after treatment or transfection. Cell lines were washed once with ice-cold PBS (1X), followed by scraping in PBS, collection into 1.7 ml microcentrifuge tubes, and centrifuged at 5,000 rpm for 2 min at 4 °C. Supernatant was aspirated, and cell pellets were frozen at -80 °C. Cell pellets were lysed and resuspended in 1X RIPA lysis buffer (Millipore Sigma) supplemented with 1X cOmplete Protease Inhibitor Cocktail (Roche) and PhosSTOP phosphatase inhibitors (Roche). Lysates were incubated on ice for 10 min and subsequently centrifuged at 14,000 x *g* for 10 min at 4 °C; the supernatant was evaluated for protein content using a Pierce BCA Protein Assay (ThermoFisher Scientific). In all experiments, 50 μg protein was loaded onto Novex™ 4-20% Tris-Glycine Mini Gels (ThermoFisher Scientific), transferred to nitrocellulose membranes with the iBlot™ 2 Gel Transfer Device (ThermoFisher Scientific), and probed with the specified antibodies overnight at 4 °C in 1X TBS-Tween (Tris-buffered saline plus 0.02% Tween 20) containing 5% non-fat milk. Immunoblots were visualized using either a LICOR Odyssey Fc imaging instrument or by development with HyBlot CL Autoradiography Film (Thomas Scientific).

### Quantitative real-time PCR (Q-PCR)

For real-time PCR experiments, total RNA was extracted from 1 × 10^6^ cells using the QIAGEN RNeasy Mini Kit (74104) following the manufacturer’s instructions. Two micrograms of total RNA were used for cDNA synthesis in a 20 μl reaction. The cDNA synthesis steps consisted first of a 5 min incubation at 65 °C of the hexameric random primers, dNTP, and RNA, followed by cycles of 25 °C for 10 min, 42 °C for 50 min, and 75 °C for 10 min with the addition of 0.1 M DTT, 5X Reaction Buffer, SuperScript III Reverse Transcriptase (18080-044), and RNaseOUT inhibitor (all from Life Technologies). The synthesized cDNA was diluted to 100 μl with molecular grade H_2_O, and quantitative PCR was conducted in 384-well plates in a 20 μl volume consisting of 2 μl diluted cDNA, 1 μl primers, 7 μl H_2_O, and 10 μl TaqMan Universal PCR Master Mix (4364340; Life Technologies). The PCR was performed using the default cycling conditions (50 °C for 2 min and 95 °C for 10 min, and 40 cycles of 95 °C for 15 s and 60 °C for 10 min) with the ABI QuantStudio 6 Flex Real-Time PCR System (Applied Biosystems).

Triplicate samples were used for the Q-PCR, and the mean values were calculated. The data in all figures represent three independent experiments. Relative gene expression was calculated from the ratio of the target gene expression to the expression of the internal reference gene (β-actin) based on the cycle threshold values.

### Transfection and siRNA knockdown

Lonza Kit V (VCA-1003) was used for electroporation of BxPC-3 cells with the Lonza Nucleofector 2b device (Cat# AAB-1001). For this line, 1 × 10^6^ cells were resuspended in the electroporation solution provided in the kit, per the manufacturer’s recommendations, and 2 µg of DNA was added to the cell suspension before transferring to a clean electroporation cuvette. This ratio of cells to DNA was maintained when scaling up experiments for immunoblot analysis. After the electric charge was applied, the cells were transferred with a transfer pipette to culture dishes containing full-serum and media. The cell-type specific program was used for the electroporation procedure. Lipofectamine RNAIMAX reagents (Thermo Fisher Scientific, Cat# 13778075) was used to transfect siRNA into CFPAC-1 cells following the manufacturer’s protocol. Lipofectamine 2000 (Thermo Fisher Scientific, Cat# 11668027) was used to transfect plasmid DNA into CFPAC-1, HT-29, and HTB134 cells. Purified E. coli genomic DNA fragments around 1000 bp in size were transfected into both BxPC-3 and CFPAC-1 cells using Lipofectamine 2000 according to the manufacturer’s protocols.

Lonza Kit V (VCA-1003) was also used for co-electroporation of siRNA and DNA into BxPC-3 cells with the Lonza Nucleofector 2b device for 48 h. 20nM of scrambled siRNA or target gene siRNAs as indicated in the figures were used for co-transfection experiments when siRNA and plasmid DNA were simultaneously co-transfected.

### ELISA

The cGAMP ELISA kit was purchased from Cayman Chemical (cat# 501700), and the manufacturer’s protocol was followed. Absorbance was quantitated at a wavelength of 450 nm using a plate reader. Triplicate studies were performed for each experimental condition, and means were determined for graphical presentation.

### Amplex Red^®^ assay to detect extracellular H_2_O_2_

The Amplex Red^®^ Hydrogen Peroxide/Peroxidase Assay Kit (cat# A22188; Life Technologies) was used to detect extracellular H_2_O_2_ generation. BxPC-3 cells were either transfected with 2 µg of pGL3-BV plasmid for 48 h or treated with IL-4 (50 ng/ml) for 24 h and then washed twice with 1 X PBS, trypsinized, and counted. 2 × 10^4^ live cells in 20 µl of 1X Krebs-Ringer phosphate glucose [KRPG] buffer was mixed with 100 µl of a solution containing 50 µM Amplex Red^®^ and 0.1 U/ml horse radish peroxidase in KRPG buffer plus 1 µM ionomycin and then incubated at 37 °C for the indicated times. The fluorescence of the oxidized 10-acetyl-3,7-dihydroxyphenoxazine was then measured at excitation and emission wavelengths of 530 nm and 590 nm, respectively, using a SpectraMax Multi-Mode Microplate Reader (Molecular Devices, Sunnyvale, CA, USA); the amount of extracellular H_2_O_2_ was calculated based on a standard curve using 0-2 µM H_2_O_2_. Each value in the figures is the mean value of quadruplicate samples.

### Quantification and statistical analysis

Data are displayed as the mean ± SD from at least triplicate experiments, unless otherwise specified. Comparisons between two groups were analyzed using the Student’s *t*-test, whereas comparisons between multiple groups were performed via ANOVA. Statistical significance was determined as a *P* value < 0.05 and shown with an asterisk (*); a *P* value < 0.01 is represented with three asterisks (***).

## Abbreviations

DUOX2: dual oxidase 2
ROS: reactive oxygen species
PDAC: pancreatic ductal adenocarcinoma
NOX: NADPH oxidase
(cGAS): cyclic GMP-AMP Synthase
(STING): Stimulator of Interferon Genes

## AUTHOR CONTRIBUTIONS

YW, SLW and JHD conceptualized the project. SLW and YW performed the experiments and analyzed data. SLW, YW, and JHD wrote the manuscript. MK, SA, JLM, GJ, JL, ID, AJ, BD, and KR edited and approved the manuscript.

## ADDITIONAL INFORMATION

### Supplementary information

The online version contains supplementary material.

**Correspondence** and requests for materials should be addressed to James H. Doroshow, M.D.

