## Supplemental Figure Legends and Figures for "Exogenous DNA upregulates DUOX2 expression and function in human pancreatic cancer cells by activating the cGAS-STING signaling pathway"

**SUPPLEMENTARY INFORMATION**

**Wang et al.**

### Supplementary Methods

#### Cell culture, antibodies, plasmids, primers, and reagents

Human pancreatic cancer cell lines: BxPC-3 (CRL-1687), CFPAC-1 (CRL-1918), AsPC-1 (CRL-1682), and Hs 766T (HTB-134); human colon cancer cell lines: Ls513 (CRL-2134), HT-29 (HTB-38), and T84 (CCL-248) were obtained from the American Type Culture Collection (ATCC, Manassas, VA). BxPC-3, AsPC-1, and Ls513 cells were cultured in RPMI-1640 medium (HyClone SH30027.01) supplemented with 10% Fetal Bovine Serum (FBS, Gemini Bio-Products 100-106) and 1% 100 mM sodium pyruvate (Gibco) by volume. Hs 766T cells were cultured in Dulbecco's Modified Eagle's Medium (DMEM) (ATCC 30-2002) supplemented with 10% FBS. CFPAC-1 cells were cultured in Iscove's Modified Dulbecco's Medium (ATCC 30-2005) supplemented with 10% FBS. HT-29 cells were cultured with McCoy's 5a Medium Modified (ATCC 30-2007) supplemented with 10% FBS. T84 cells were cultured with DMEM:F-12 Medium (ATCC 30-2006) supplemented with 5% FBS.

Antibodies against cGAS (15102), STING (3337), p-NF-κB $p65_{S536}$ (3033), NF-κB p65 (8242), TBK1 (38066), p-TBK1 (5483), p-$IRF3_{S396}$ (4947), IRF3 (4302), p-$Stat2_{Y690}$ (88410), IRF9 (76684), HIF1β (3718), Stat1 (9175), p-$Stat1_{Y701}$ (9167), γ-$H2AX_{S139}$ (2577), Lamin A/C (2032), and β-actin (3700) were purchased from Cell Signaling Technologies (CST, Beverly, MA). HIF-1α (610959) and Stat2 (610187) antibodies were from BD transduction Laboratory. IRF1 (s497) antibody was purchased from Santa Cruz Biotechnology (Santa Cruz, CA). Mouse monoclonal antibody, which reacts with both human DUOX1 and DUOX2, was developed by Creative Biolabs (Port Jefferson Station, NY) and characterized by our laboratory [1].

The plasmids used in these studies included: pGL3-Basic Vector (pGL3-BV, E1751), Bacterial strain JM109 for bacterial genomic DNA extraction (P9751) from Promega, pcDNA3.1-HA plasmid (pcDNA3,128034) from Addgene, and pReceiver-M08 plasmid(EX-NEG-M08) from GeneCopoeia. The STING agonist MSA-2 (HY-136927) was from MedChem Express. Recombinant human cytokines Human IL-4 (204-IL-050), Human IL-17A (317-ILB-050), Human IFN-α (11100-1) and Human IFN-β (8499-IF-010) were from R & D Systems. 2'-3' cGAMP (tlrl-nacga23-5) was obtained from InvivoGen,

The following primers used for Q-PCR in this study were purchased from Applied Biosystems: DUOX2 (Hs00204187_m1), DUOXA2 (Hs01595310_m1), DUOX1 (Hs00213694_ml), DUOXA1 (Hs00328806_m1), NOX1 (Hs00246589_ml), cGAS (Hs00403553_m1), STING (Hs00736955_g1), IRF3 ( Hs01547283_m1), IRF1 (Hs00971965_m1), β-actin (Hs01060665_g1), IFN-β (Hs01077958_s1), IRF-9 (Hs00196051_ml), STAT2 (Hs01013119_g1), RELA ( Hs01042010_ml), and STAT1 (Hs00234829_m1). IRF-1 siRNA-A: Silencer Select Human IRF-1 siRNA, was from Ambion (s7501); IRF-1 siRNA-B: On-Target plus Smart pool human IRF1, was from Dharmacon (011704-00-10). RELA siRNA, On-Target plus SMART pool Human RELA (L-003533-00) was obtained from Dharmacon; RELA-A and B, RELA Silencer select siRNA, were from Ambion (S11914 and S11915); On-Target plus Human STAT1 siRNA, Smart pool, was purchased from Dharmacon (L-003543-00-0010). cGAS siRNA-2, On-Target plus Human MB21D1 siRNA, SMART Pool, was from Dharmacon (L-015607-02-0020); cGAS siRNA-3, Silencer pre-designed siRNA, was from Ambion (s129126). On-Target plus Human STAT2 siRNA-A, Smart Pool(L-012064-00-0020); On-Target plus SMART siRNA Human STAT2-B (J-012064-08-0010); and On-Target plus SMART siRNA Human STAT2-C (J-012064-06-0010) were from Dharmacon. IRF3 siRNA-A and B, IRF3 Silencer Select siRNA (s7508 and s7509) were from Ambion; IRF3 siRNA-C: siGENOME human IRF3 siRNA SMART pool (M-006875-02-0020) was from Dharmacon.

### Supplementary Figure Legends

**Supplementary Fig. S1 Effect of exogenous DNA or proinflammatory cytokines on NADPH oxidase expression in human colon cancer cell lines.** **A** Ls513 human colon cancer cells were transfected with either two different DNA plasmids or treated with IL-17A and examined for the expression of NOX1 or DUOX2 48 h after transfection or 24 h following cytokine treatment, respectively. *$P < 0.05$. **B** DUOX2 expression was examined in T84 human colon cancer cells 48 h following transfection with a DNA plasmid or after a 24 h exposure to the combination of IL-17A and IL-4 evaluated as a positive control. *$P < 0.05$. **C** DUOX2 mRNA expression was determined by RT-PCR in HT-29 human colon cancer cells 48 and 72 h following transfection with the pGL3-BV plasmid or exposure to the combination of IL-17A and IL-6 for 72 h. *$P<0.05$. These data represent results from three independent experiments.

**Supplementary Fig. S2 Concentration- and time-dependent effects of the STING agonist MSA-2 and cGAMP on DUOX expression and cGAS-STING signal transduction in human pancreatic cancer cells.** **A** Comparison of the effects of two concentrations of the STING agonist MSA-2 versus IL-17A on the expression of DUOX2 in BxPC-3 cells. Tumor cells were exposed to MSA-2 for 48 h and to IL-17A for 24 h. *$P < 0.05$. **B** Relationship of MSA-2 concentration to the expression of DUOX2 following 48 h of drug exposure in CFPAC-1 pancreatic cancer cells. *$P < 0.05$. **C** Time course examining the relationship between time of exposure for cGAMP and MSA-2 and DUOX expression and activation of cGAS-STING signal transduction in the BxPC-3 cell line. **D** Relationship between time of exposure for cGAMP and MSA-2 as well as co-treatment with dexamethasone on DUOX expression, cGAS-STING signaling, and DNA double strand scission in CPFAC-1 cells. The results presented represent at least three independent experiments.

**Supplementary Fig. S3 Role of Stat1 and Stat2 in plasmid-enhanced DUOX2 mRNA expression.** **A** Effect of Stat1 siRNA on pGL3-BV-enhanced expression of DUOX2 (right panel) as well as on Stat1 expression itself (left panel) in BxPC-3 cells. Real time PCR for DUOX2 and Stat1 mRNA was performed 48 h following plasmid and siRNA co-transfection. ***$P < 0.01$. **B** Effect of Stat2 siRNA on the expression of DUOX2 (left panel) or Stat2 (right panel) following pGL3-BV transfection in BxPC-3 cells; exposure times for plasmid and siRNA were identical to Supplementary Fig. S3A. ***$P < 0.01$. **C** Effect of Stat2 siRNA on the expression of DUOX2 (left panel) and Stat2 (right panel) 24 h after siRNA transfection prior to a 24 h exposure to IFN-β or a 24 h treatment with MSA-2 (10 μM). *$P < 0.05$. **D** Effect of multiple Stat2 siRNAs on DUOX2 (left panel) or Stat2 (right panel) expression following IFN-β treatment. Experiments performed as in figure S3C. ***$P < 0.01$. **E** Effect of multiple Stat2 siRNAs on DUOX2 (left panel) or Stat2 (right panel) expression following pGL3-BV transfection. QPCR for DUOX2 and Stat2 mRNA was performed 48 h following plasmid and siRNA co-transfection. ***$P < 0.01$. **F** Effect of multiple Stat2 siRNAs on DUOX2 (left panel) or Stat2 (right panel) expression following cGAMP treatment. Stat2 siRNAs transfected into BxPC-3 cells 24 h before initiation of a 24 h exposure to cGAMP. ***$P < 0.01$. **G** Effect of IRF1 siRNAs on plasmid-enhanced DUOX2 mRNA expression. Real time PCR for DUOX2 and IRF1 mRNA was performed 48 h following plasmid and siRNA co-transfection. ***$P < 0.01$. These studies were conducted in triplicate.

### Suppl. Fig. S1

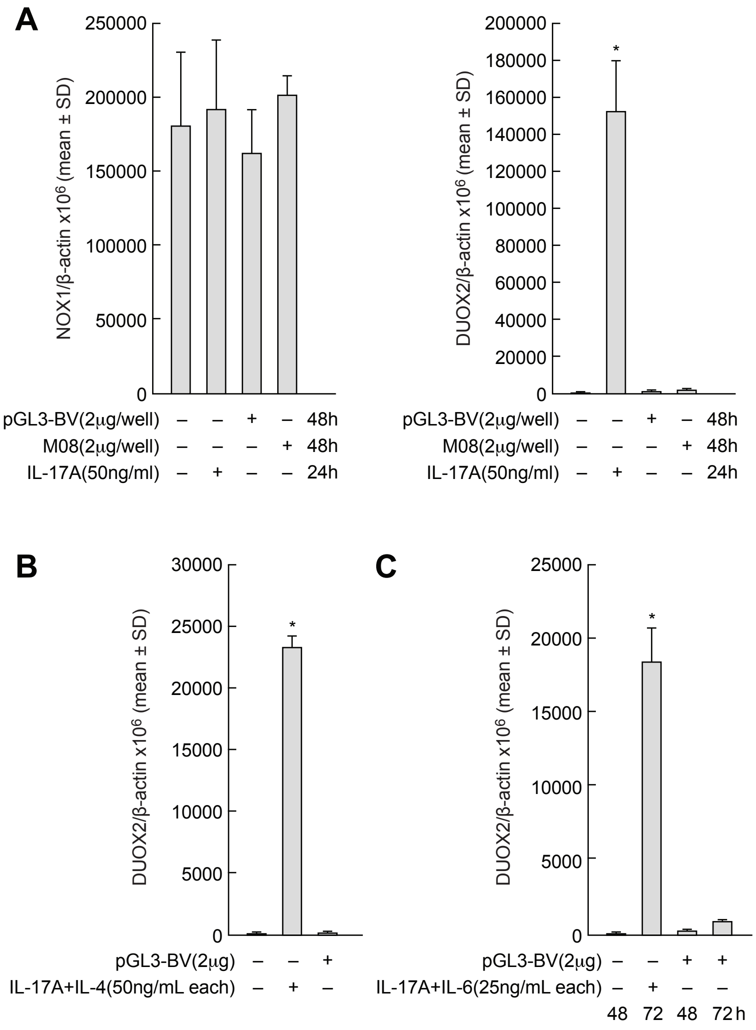

A
NOX1/β-actin x10^6 (mean ± SD)
250000
200000
150000
100000
50000
0
pGL3-BV(2μg/well) – – + – 48h
M08(2μg/well) – – – + 48h
IL-17A(50ng/ml) – + – – 24h
DUOX2/β-actin x10^6 (mean ± SD)
200000
180000
160000
140000
120000
100000
80000
60000
40000
20000
0
*
pGL3-BV(2μg/well) – – + – 48h
M08(2μg/well) – – – + 48h
IL-17A(50ng/ml) – + – – 24h
B
DUOX2/β-actin x10^6 (mean ± SD)
30000
25000
20000
15000
10000
5000
0
*
pGL3-BV(2μg) – – +
IL-17A+IL-4(50ng/mL each) – + –
C
DUOX2/β-actin x10^6 (mean ± SD)
25000
20000
15000
10000
5000
0
*
pGL3-BV(2μg) – – + +
IL-17A+IL-6(25ng/mL each) – + – –
48 72 48 72 h

### Suppl. Fig. S2

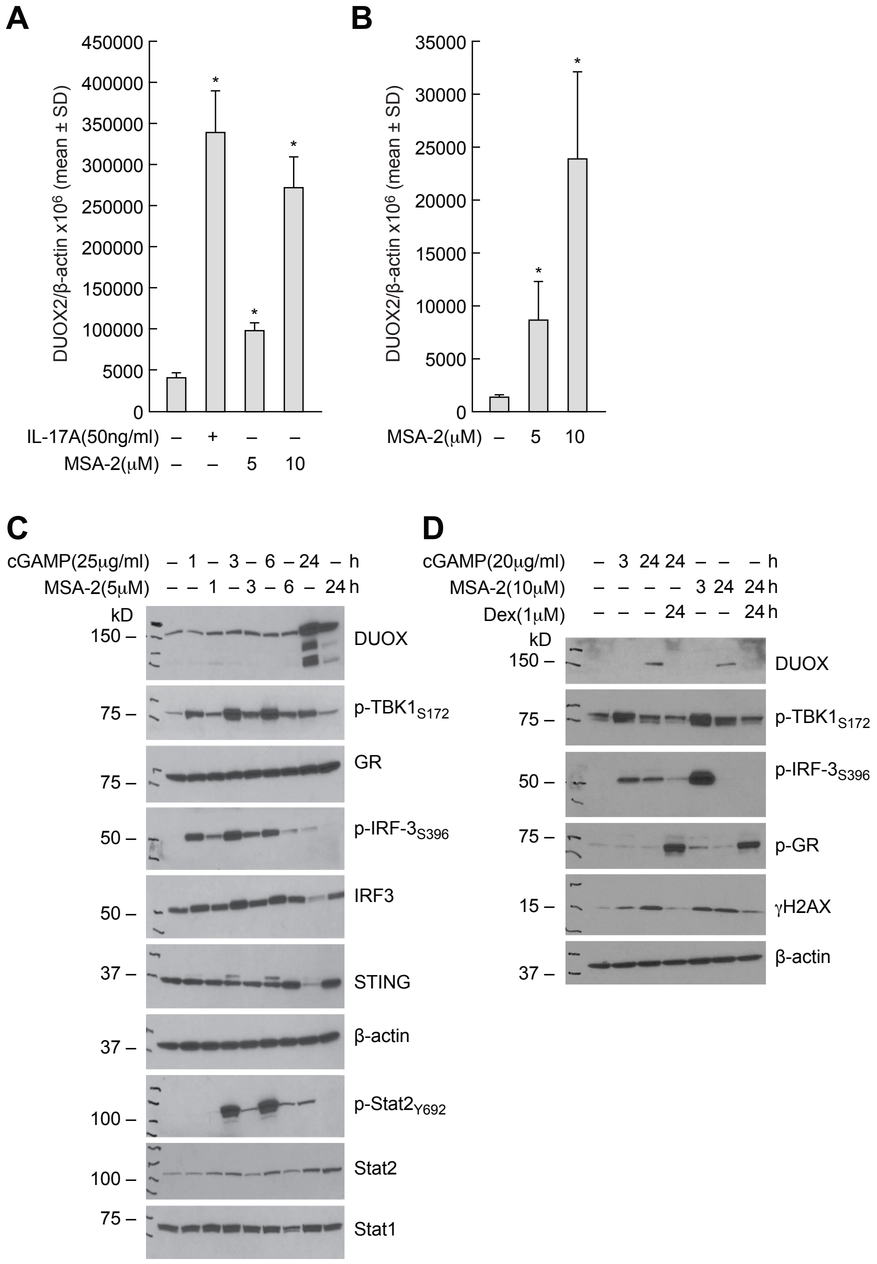
A
DUOX2/β-actin x10^6 (mean ± SD)
450000
400000
350000
300000
250000
200000
150000
100000
5000
0
IL-17A(50ng/ml) – + – –
MSA-2(μM) – – 5 10
B
DUOX2/β-actin x10^6 (mean ± SD)
35000
30000
25000
20000
15000
10000
5000
0
MSA-2(μM) – 5 10
C
cGAMP(25μg/ml) – 1 – 3 – 6 – 24 – h
MSA-2(5μM) – – 1 – 3 – 6 – 24 h
kD
150 –
DUOX
75 –
p-TBK1S172
GR
75 –
50 –
p-IRF-3S396
IRF3
50 –
37 –
STING
β-actin
37 –
p-Stat2Y692
100 –
Stat2
100 –
75 –
Stat1
D
cGAMP(20μg/ml) – 3 24 24 – – – h
MSA-2(10μM) – – – – 3 24 24 h
Dex(1μM) – – – 24 – – 24 h
kD
150 –
DUOX
75 –
p-TBK1S172
50 –
p-IRF-3S396
75 –
p-GR
15 –
γH2AX
β-actin
37 –

**Suppl. Fig. S3**

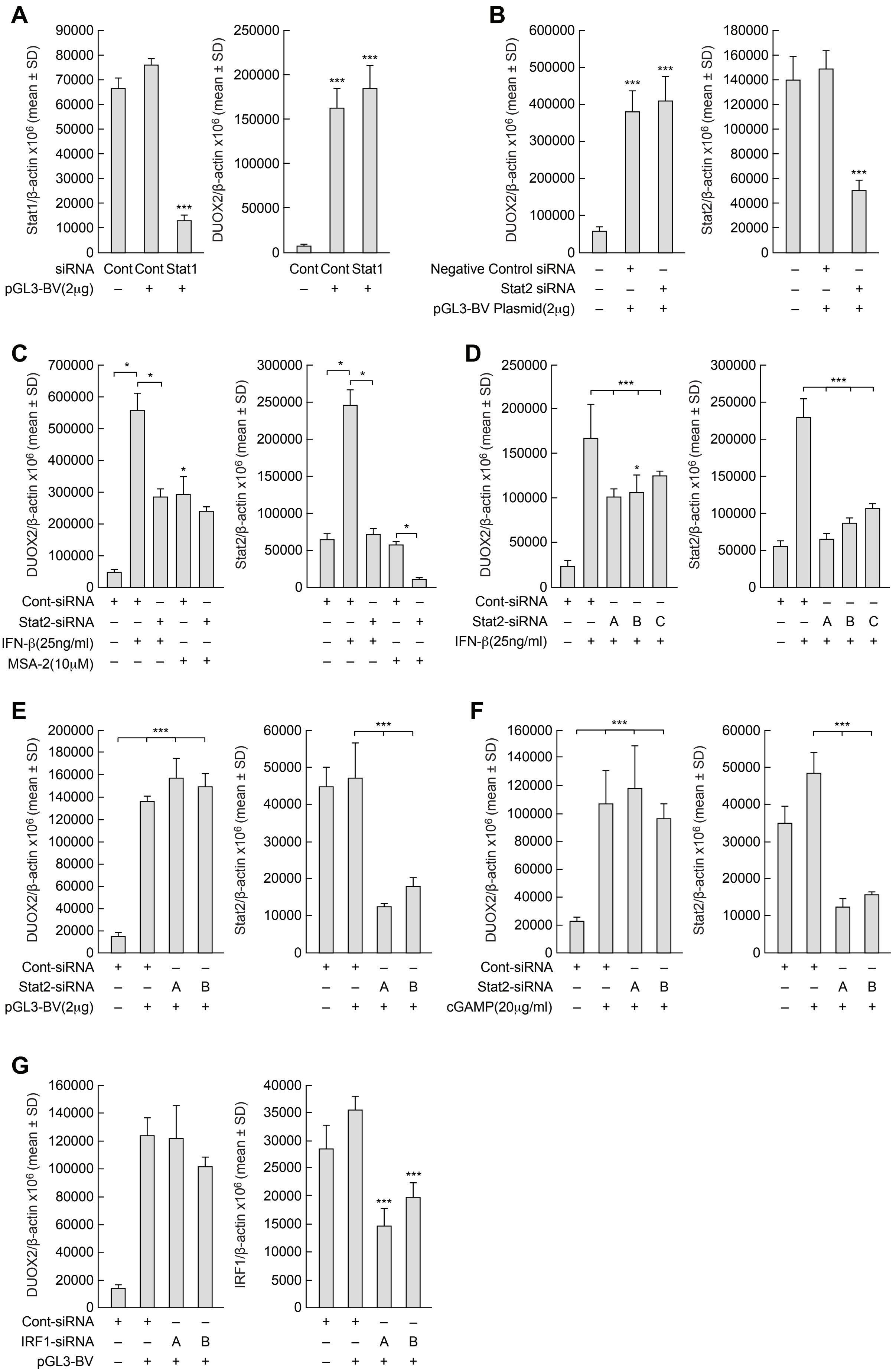

A
Stat1/β-actin x10⁶ (mean ± SD)
DUOX2/β-actin x10⁶ (mean ± SD)
siRNA Cont Cont Stat1
pGL3-BV(2μg) – + +
***
B
DUOX2/β-actin x10⁶ (mean ± SD)
Stat2/β-actin x10⁶ (mean ± SD)
Negative Control siRNA
Stat2 siRNA
pGL3-BV Plasmid(2μg)
C
DUOX2/β-actin x10⁶ (mean ± SD)
Stat2/β-actin x10⁶ (mean ± SD)
Cont-siRNA
Stat2-siRNA
IFN-β(25ng/ml)
MSA-2(10μM)
D
DUOX2/β-actin x10⁶ (mean ± SD)
Stat2/β-actin x10⁶ (mean ± SD)
Cont-siRNA
Stat2-siRNA – – A B C
IFN-β(25ng/ml)
E
DUOX2/β-actin x10⁶ (mean ± SD)
Stat2/β-actin x10⁶ (mean ± SD)
Cont-siRNA
Stat2-siRNA – – A B
pGL3-BV(2μg)
F
DUOX2/β-actin x10⁶ (mean ± SD)
Stat2/β-actin x10⁶ (mean ± SD)
Cont-siRNA
Stat2-siRNA – – A B
cGAMP(20μg/ml)
G
DUOX2/β-actin x10⁶ (mean ± SD)
IRF1/β-actin x10⁶ (mean ± SD)
Cont-siRNA
IRF1-siRNA – – A B
pGL3-BV
